# Spatial frequency representation in V2 and V4 of macaque monkey

**DOI:** 10.1101/2022.07.27.501743

**Authors:** Ying Zhang, Kenneth E. Schriver, Jia Ming Hu, Anna Wang Roe

## Abstract

Spatial frequency (SF) is an important attribute in the visual scene and is a defining feature of visual processing channels. However there remains many unsolved questions about how primate visual cortex, in particular extrastriate areas V2 and V4, codes this fundamental information. Here, using intrinsic signal optical imaging in visual cortex of Macaque monkeys, we quantify the relationship between spatial frequency maps and (1) visual topography, (2) color and orientation maps, and (3) across visual areas V1, V2, and V4. We find that in orientation regions, low to high spatial frequency is mapped orthogonally to orientation; however, in color regions, which is reported to contain orthogonal axes of color and lightness, only low spatial frequencies are represented without a gradient of SF representation. This produces the first observation of a population spatial frequency fluctuation related to the repeating color/orientation organizations. These findings support a generalized hypercolumn model across cortical areas, comprised of two orthogonal parameters with additional parameters.

## Introduction

Spatial frequency (SF) selectivity is a fundamental feature encoded in the visual system. Previous studies have shown that the organizations of SF selectivity are related to orientation and color maps in primary visual cortex (V1), and have a high degree of periodicity in both cats (Hübener et al., 1997; Issa et al., 2000; Shoham et al., 1997; Tootell et al., 1981) and monkeys (Silverman et al., 1989). Studies have consistently shown an orthogonal mapping of spatial frequency and orientation, suggesting an efficient arrangement that provides each orientation access to a wide range of spatial frequencies (Issa et al., 2000; Nauhaus et al., 2012, 2016; Xu et al., 2007). In contrast, color representation in V1 (blobs) is generally associated with a range of lower spatial frequencies (Silverman et al., 1989; Tootell et al., 1988). Thus, while there is a high to low gradient of SF representation across eccentricities, within this topography there are further SF organizations. This systematic architecture in V1 suggests that spatial frequency may be a fundamental feature of the cortical ‘hypercolumn’ (c.f., Silverman et al., 1989: organized cortical modules; Hübener et al., 1997: ‘mosaics’ of functional domains for the different properties; Swindale et al., 2000: uniform coverage of cortical maps).

Whether there are systematic associations between SF and other parameters in extratriate areas, such as V2 and V4, is not known. The traditional view of the V2 hypercolumn comprises the alternating thin-pale-thick-pale stripe cycle (Roe and Ts’o, 1995). Within thin stripes, surface properties, typically associated with low SF preferences, such as hue maps (Xiao et al., 2003), ‘brightness’ maps (Roe et al., 2005), and ON/FF maps (Wang et al., 2007) are represented. Within the thick and pale stripes are higher order orientation maps such as those defined by illusory contours (Ramsden et al., 2001), motion direction maps (Lu et al., 2010) and maps for motion-defined edges (Chen et al., 2016), as well as stereo-defined Near-to-Far disparity maps (Chen et al., 2008). Neuronal tuning for other features such as texture have also been described (Freeman et al., 2013), but functional organization has not yet been investigated. However, there is little systematic data relating SF representation in V2 to functional stripes (cf. Gegenfurtner et al., 1996; Levitt et al., 1994; Tootell and Hamilton, 1989) and, despite previous attempts, few study has demonstrated functional mapping of stripes based on SF alone (Lu and Roe, 2007, Lu et al., 2018).

In V4, surface and shape information is organized into, for lack of better terminology, the ‘color’ and ‘orientation’ bands. Within the color bands, maps for hue and for luminance have been desribed (Tanigawa et al., 2010; Liu et al., 2020; Li et al., 2022); within orientation bands are maps for contrast-defined contours (Hu et al., 2020; Li et al., 2013; Lu et al., 2018; Tang et al., 2020; Tanigawa et al., 2010), disparity-defined contours (Fang et al., 2019), as well as maps for curvature degree and curvature orientation (Hu et al., 2020; Ponce et al., 2017). Despite our growing understanding of functional organization in V4, how SF preference maps (first reported in Lu et al., 2018) coordinate with other feature maps in V4 remains unknown.

As part of our investigation into what are ‘hypercolumns’ in extrastriate cortical areas, we propose a general hypercolumn layout for V2 and V4 that includes SF (cf. Roe et al., 2009; Ts’o et al., 2009). We predict that in each area (1) the range of spatial frequency associations shift with the topographic location of the ‘hypercolumn’ (Figure 1A), (2) orientation selective regions (Figure 1B, blue) have a range of low (light gray) to high (dark gray) SF representation; iso-SF contours (blue dashed lines) map orthogonally to iso-orientation contours (red dashed lines), and (3) color selective regions (Figure 1B, orange) exhibit an association with a range of low spatial frequencies (light gray). To address this proposal, using intrinsic signal optical imaging (ISOI) in Macaque monkey, we imaged large cortical fields of view that contained sufficient territory to allow comparisons of functional organization across eccentricities. Quantification of the relationship between SF maps and color and orientation maps in V2 and V4 revealed organizations that generally support our proposal for a hypercolumn architecture.

**Figure 1.**
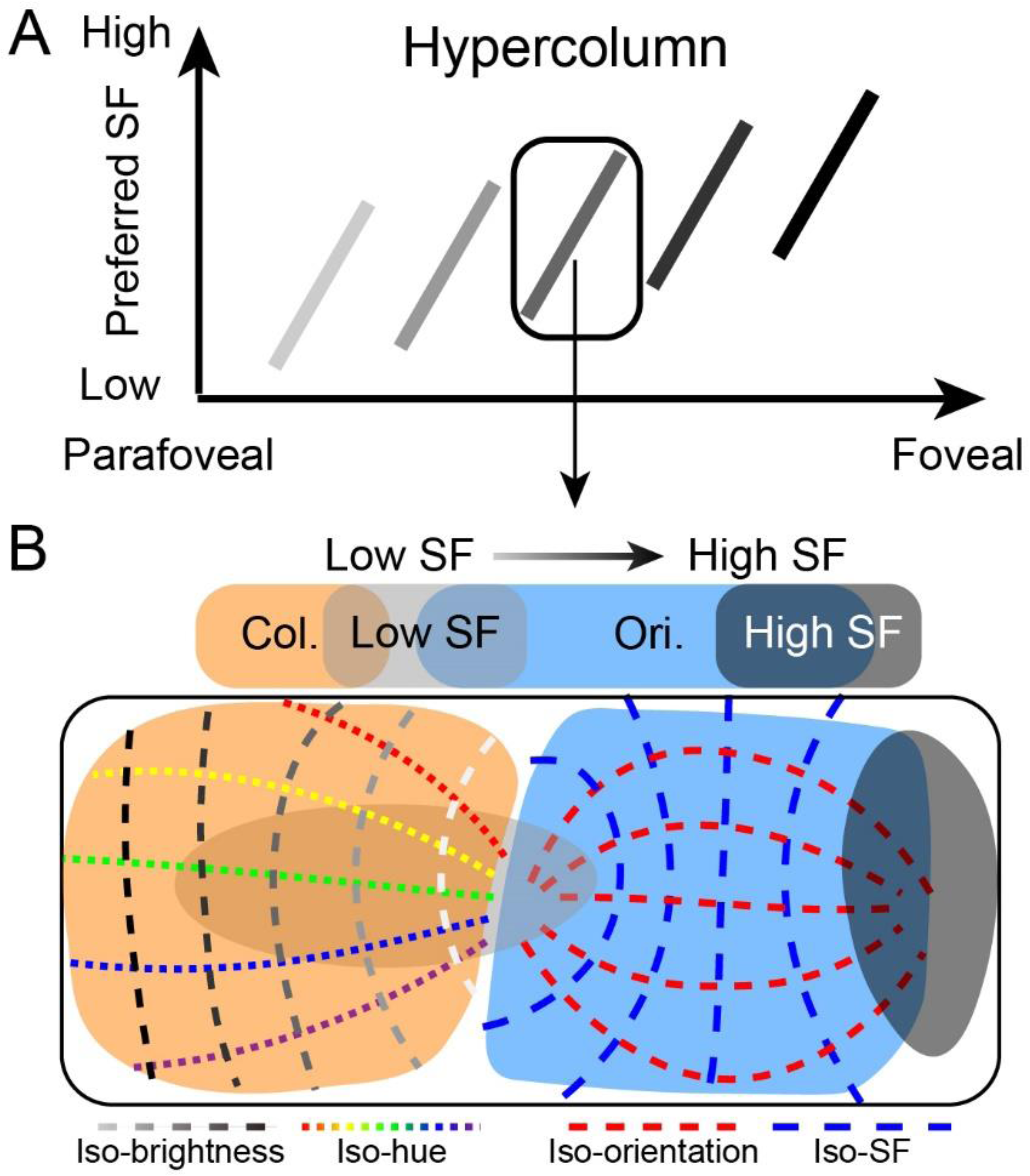
Illustration of hypercolumn (including SF, color and orientation domains) in the visual cortex. A. As eccentricity decreases from parafoveal to foveal region, the preferred SF gradually increases (representing as the brightness of the short bar decreases). However, for a local region (marked by a rectangle), it covers a full range of SF representations in its corresponding topographic locations. This local region can be considered as a ‘hypercolumn’. B. Details of structure in a single hypercolumn. In this local region, color domains (orange) and orientation domains (cyan) exhibit different relationships with SF domains (low SF preference domain: light region; high SF preference domain: dark region). Orientation maps orthogonally to spatial frequency (see iso-orientation contours and iso-SF contours), and a large range of spatial frequencies are available to each orientation. In comparison, color domains tend to have more spatial overlap with low SF preference domains, while avoid overlapping with high SF preference domains. In color domains, there exists another orthogonal relationship between hue (colored dotted lines: iso-hue contours) and brightness (dotted lines in different darknesses: iso-brightness contours).

## Results

### Overall SF preference across imaged visual cortical areas

Imaging a large field of view of the cortex enables us to directly compare the response differences among different visual areas. Figure 3 shows the blood vessel maps (3A, 3D) and corresponding SF preference maps (3B, 3E) for two separate cases wherein we imaged regions spanning V1, V2, and V4. The SF preference maps (Figure 3B, E) were generated by calculating the response amplitude of each pixel in these areas for stimuli with six different SFs, each at orientations of 45° and 135° . We found that in both cases, most of the imaged V1 region favored high SF stimuli, while most of the imaged V4 region favored low SF stimuli. To further illustrate the distribution of SF preference in each area, we calculated a coverage ratio for each SF. For V1, the coverage ratio peaks at high SF (light gray bars in Figure 3C and F), while for V4, the coverage ratio peaks at low SF (black bars in Figure 3C and F), which is consistent with previous findings that V4 prefers lower SF than V1 (Lu et al., 2018). Our maps, in contrast to earlier studies, directly show the overall SF preference in cortical space.

### Shift of selective activation center according to SF

Although several studies have shown that gratings with different SFs produce different response patterns (Hübener et al., 1997; Lu and Roe, 2007; Lu et al., 2018; Shoham et al., 1997), surprisingly, most of them placed a strong emphasis on response amplitude modulation, with little quantification of the spatial distribution features of the activated network. Figure 4 illustrates the spatial distribution of functional domains and the shift in the preferred SF across the imaged region of V4. Large scale imaging allowed us to capture highly structured maps of functional domains (e.g. differential orientation maps in Figures 4A and S1A) and reveal changes to those maps for different SF conditions. In the imaging results shown in Figure 4A, we acquired differential orientation maps using six different SFs. We found that the functional map at a specific SF is not always easily ascertained. Gratings with low SFs evoked clear selective responses (Figure 4A, right panel) in all imaged V4 regions, whereas gratings with high SFs only evoked responses in certain regions, mostly in the lateral part of cortex. We analyzed the spatial distribution of the selectively activated domains (see Figure 4A right panel), as these activated domains represent the functioning cortical regions (able to distinguish 45° vs 135° orientations) for the tested SF. For each orientation map obtained at a given SF, we defined the ‘selective activation center’ as the geometric centroid of all significantly activated pixels (two tailed t test, p<0.01). As SF increases, the location of the selective activation center shifts from medial to lateral across the cortex (Figure 4C, blue dots).

By using sets of moving gratings in two orthogonal orientations at three different SFs, we confirmed this spatial shift in activation for additional cases (Figure 4-figure supplement 1, 4 cases from 4 hemispheres of 3 monkeys). We found again that orientation selective response is spread widely over V4 under low SF conditions, whereas the response was mostly observed in the lateral area under high SF conditions. The lack of response in the medial region under high SF conditions leads to a spatial shift of the selective activation center from medial to lateral as SF is increased (Figure 4-figure supplement 1B).

The exposed V1s are in the lateral region of the hemisphere, corresponding to eccentricity of 0-2 degrees; here low SFs barely evoke measurable selective orientation responses (Figure 4-figure supplement 1A left column, Cases 1-3). The orientation map in the lateral region is only apparent at high SFs (Figure 4- figure supplement 1A, >1 cycle/deg.). Although the optimal SF differs between V1 and V4, for both areas the selective activation center moves from medial to lateral as SF increases (Figure 4-figure supplement 1, Cases 1-4, dots of different shading correspond to the activation centers at different SFs). In the monkey visual cortex, the foveal regions of V1, V2, V4 locate in the lateral part of the cortex (Gattass et al., 1981; Gattass et al., 1988, and see Figure 2D), thus our results are in agreement with the findings from electrophysiology studies that foveal regions favor high SF and play an important part in processing visual information that requires high spatial resolution.

**Figure 2.**
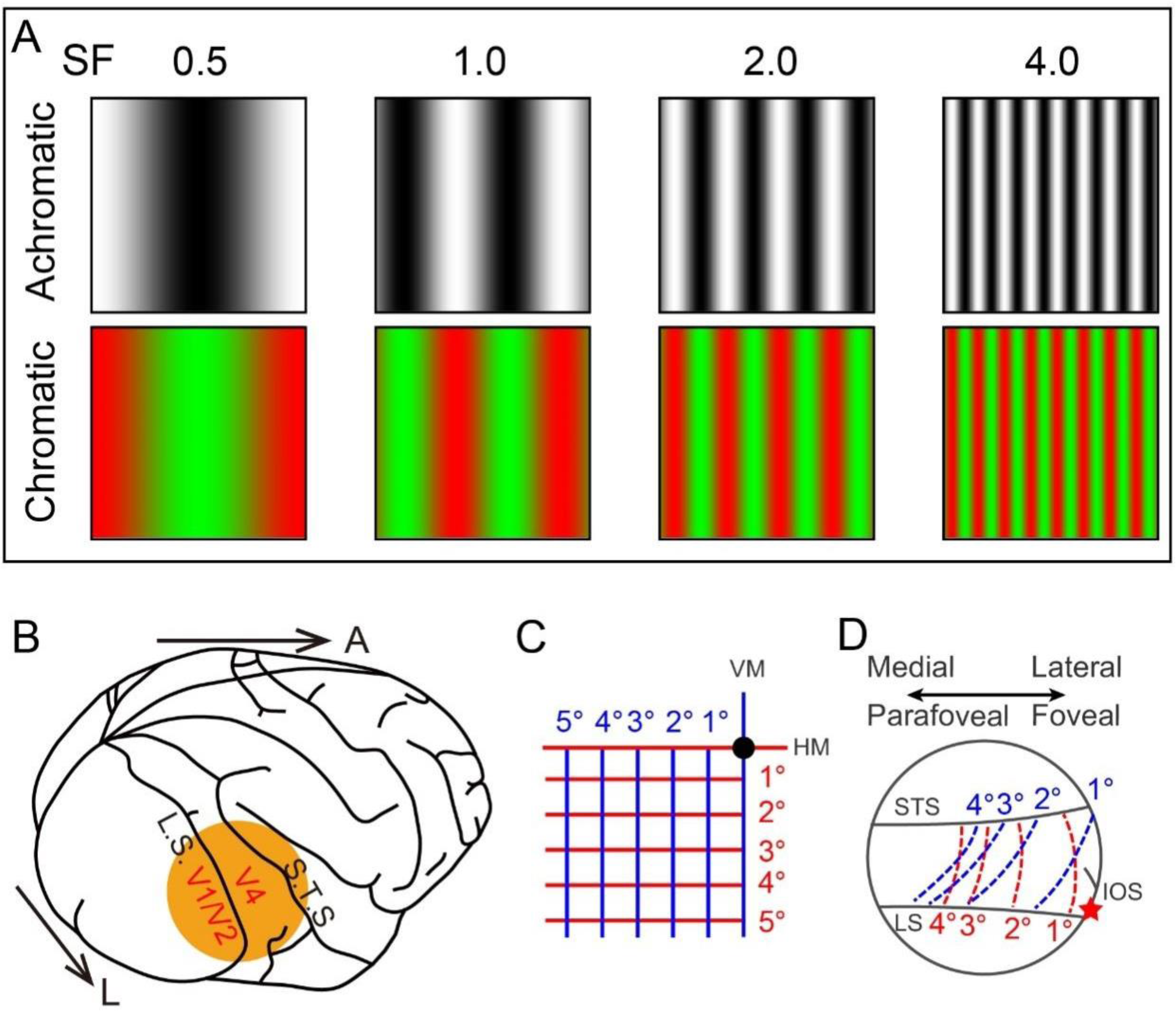
**Experimental Parameters.** A. Visual stimuli gratings. Top and bottom rows show the black/white and green/red sinusoidal gratings for four different spatial frequencies (indicated by the numbers on top; units of cycles/degree) respectively. For illustration, the stimulus size is set to 2 degrees. B. Diagram of imaging site on the cortex. L: lateral; A: anterior. C. Lower left visual field, black dot indicates fovea. Horizontal and vertical lines are colored in red and blue, respectively. D. Plot of the estimated activation locations corresponding to different lines in (C) on the imaged V4 region (corresponding to the orange disc in B). The lateral part of the imaged region corresponds to the foveal region, while the medial part corresponds to the parafoveal region. LS: lunate sulcus; STS: superior temporal sulcus; IOS: inferior occipital sulcus. Red star: estimated foveal location.

Many previous V4 imaging studies also employed grating stimuli to obtain color maps, however most of these studies (Li et al., 2014; Tanigawa et al., 2010) have not addressed how SF affects color selective response. To test this, we recorded cortical responses to red/green isoluminance sinusoidal gratings with six different SFs (Figure 4B) and compared the responses between white/black and red/green stimuli with the same SF to generate color maps. Color domains that showed significantly stronger response (two detailed t test, p<0.01, N=30) to red/green stimuli were marked by white (Figure 4B right panel). Similar to orientation domains, under low SF conditions (<1 cycle/deg.), color domains are detected in the medial region of the cortex. At high SF, the color selective response is no longer easily discernible in the medial region (SF=3, 4 cycles/deg., Figure 4B right panel). But in the lateral cortical region, color selective response was detected regardless of SF. Thus, the geometric centroid of color selective response also shifted from medial to lateral as SF increases (Figure 4C, orange dots). In a second case, we obtained similar results that different SFs evoked distinct color selective response patterns (Figure 4-figure supplement 2A) and the activation center shifts towards lateral as SF increased (Figure 4-figure supplement 2B).

In addition, we find that the coverage of the orientation and color domains appear to be underestimated when only a few spatial frequencies are tested (see numbers in Figure 4A, B and Figure 4D, E).

To further confirm the spatial relationship of functional maps acquired at different SFs, we calculated the correlation value of these functional maps. Each response map was divided into two halves: foveal region, the left half close to V4 foveal region; and parafoveal region, the right half away from foveal region. The correlation values of the two halves were calculated separately (Figure 4-figure supplement 3A and B, left matrix: foveal region; right matrix: parafoveal region). For orientation functional maps (Figure 4-figure supplement 3A), high correlation values (>0.5) appeared in the comparisons among high SF conditions in foveal region. In contrast, high correlation values appeared in the comparisons among low SF conditions in parafoveal region. For color functional map (Figure 4-figure supplement 3B), higher correlation values were measured under low SF conditions in both foveal and parafoveal regions, while high correlation values appeared only in the foveal region under high SF conditions.

These differences reflect distinct capabilities of different cortical regions (fovea vs. parafovea). Independent of the type of visual information presented (orientation or color), parafoveal regions tend to process the visual input containing low SF components, while visual input containing high SF components are better processed in foveal regions. In V4, the foveal region is capable of processing visual information with a broad range of SF sensitivity.

### Relationship between SF and orientation maps in V4 and V2

Having obtained orientation- and SF-preference maps from the same region of cortex, it becomes possible to analyze the spatial relationships between these maps. We chose V4 regions which showed strong orientation selective responses (e.g. regions marked by black dashed lines in Figure 3B, with corresponding orientation maps shown in Figure 4A), and determined the iso-orientation and iso-SF contours based on the smoothed orientation angle preference map and SF preference map (see Figure 5A and B). In both comparisons, the iso-orientation and iso-SF contours predominantly intersect orthogonally (see Figure 5C). To quantify this observation, we calculated the intersection angle of the two map gradients (orthogonal to the contours) at each pixel. Note that we calculated the angle between two gradient axes and chose the intersection angle which was smaller than or equal to 90°. The foremost intersection angle for the selected V4 region in Figure 5D falls between 80° and 90°. In the same case, V2 was also imaged (see Figure 3B). Using the same method as for V4, we calculated the intersection angle of the SF and orientation preference map gradients at each pixel. The predominant intersection angle in V2 was also near 90° (Figure 5H). Figure 5-figure supplement 1 shows additional V2 and V4 imaging regions that also had intersection angles around 90°.

**Figure 3.**
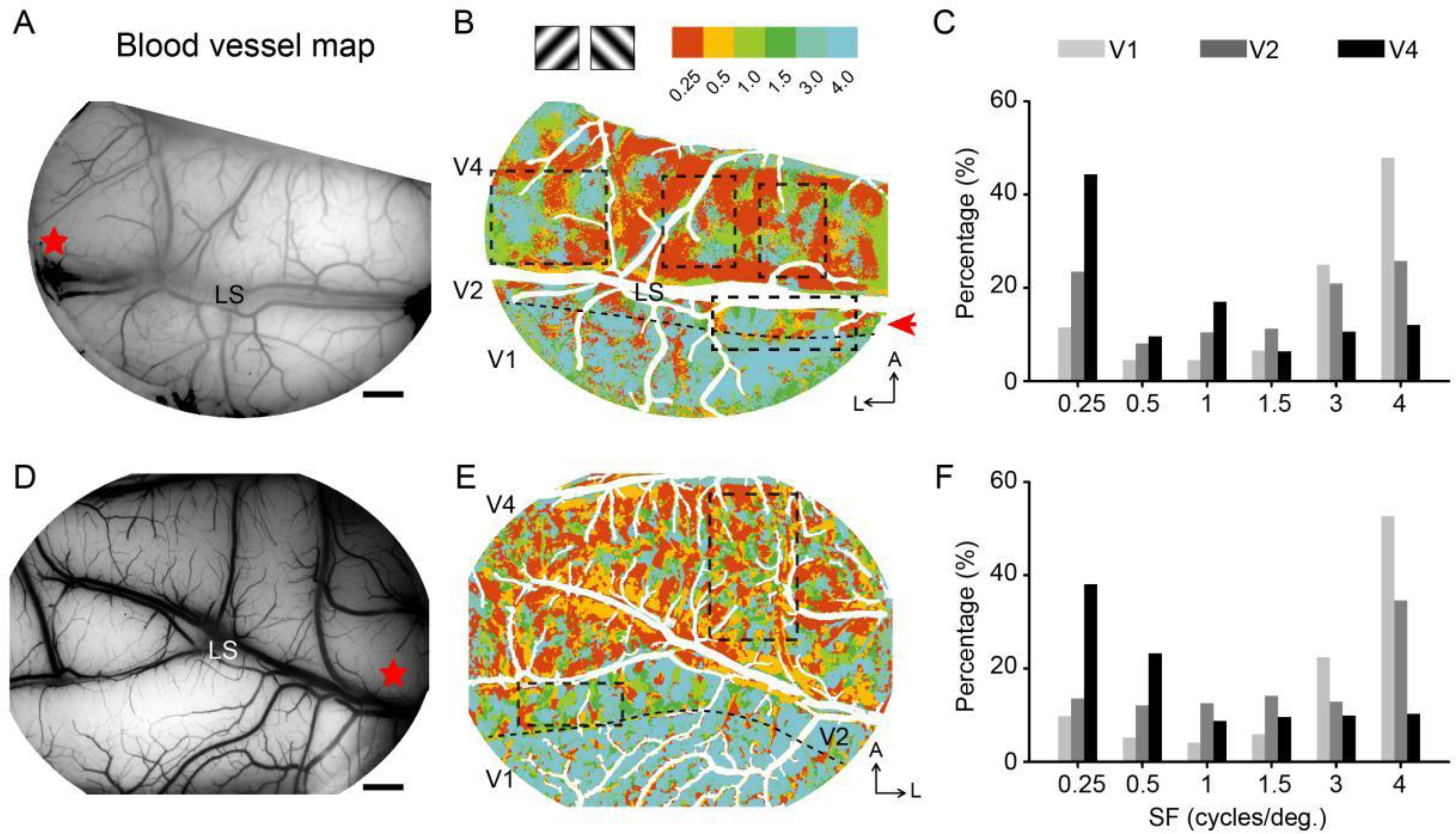
**Two examples of overall SF preference in visual cortex.** A, D: Blood vessel map of the imaged region for Case 1 and Case 2, respectively. V2 and V4 are separated by the Lunate sulcus (LS). Scale bar: 2 mm. Red star: estimated foveal location. B, E: SF preference maps for Case 1 and 2, respectively. Each SF stimulus contains two orientations, 45°and 135°. For each pixel, the preferred SF is defined as the SF corresponding to its strongest response (amplitude averaged from frame 10 to frame 20 in each condition). Different colors represent different SF preferences (see color bar at top). The border between V1 and V2 is indicated by a black dashed line. A, anterior; L, lateral. C, F. The coverage ratio of SF preference in each visual cortical area (light gray: V1; medium gray: V2; black: V4).

**Figure 4.**
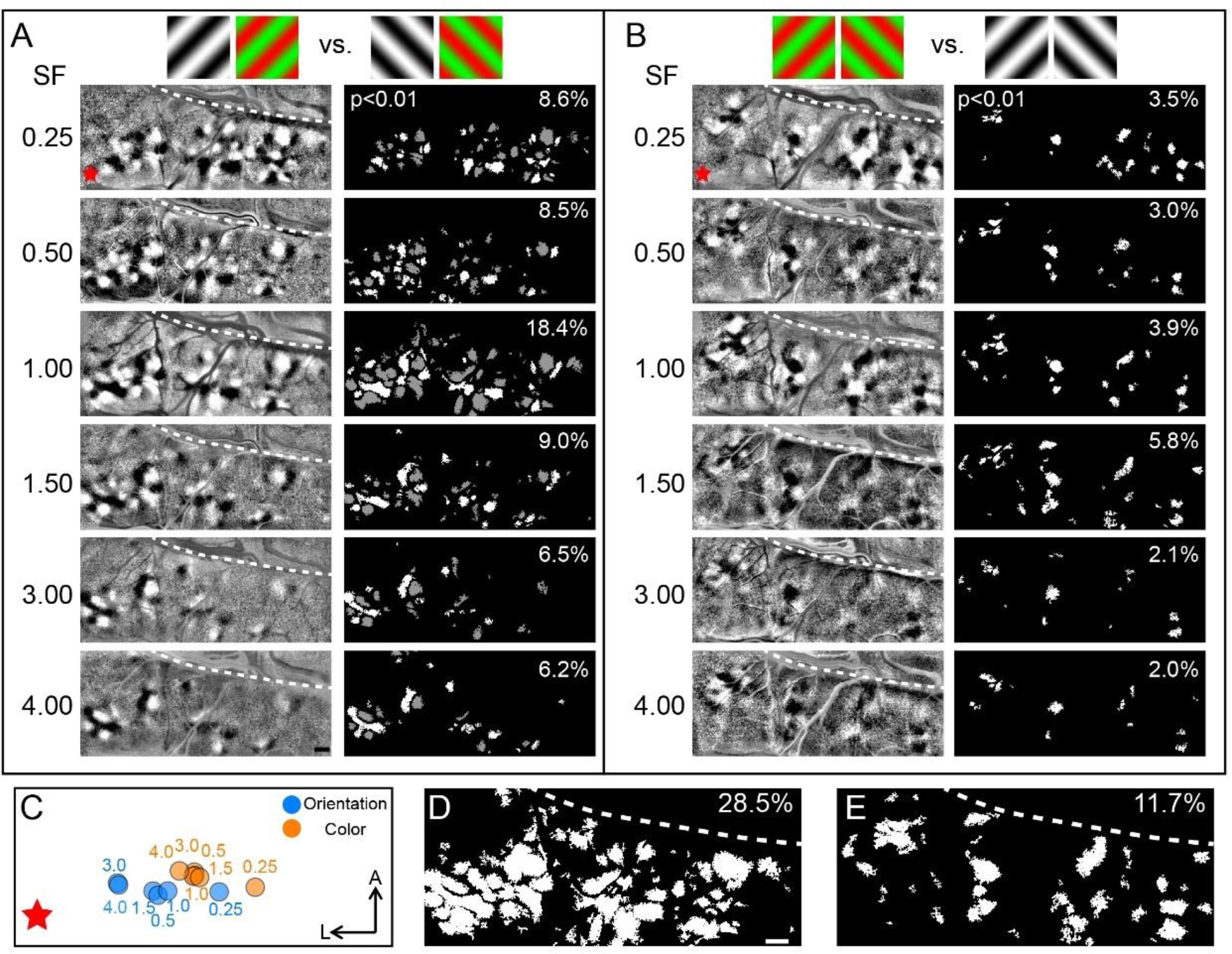
**Comparison of the functional maps obtained at different SFs.** A. V4 orientation maps and corresponding activated regions for different spatial frequencies. Gratings above the maps indicate the subtraction pair for the maps. Left panel, differential maps in response to 45° (black patches) versus 135° (white patches); right panel, stimuli-activated orientation selective regions, only including pixels that can distinguish 45°(white pixels) from 135°(gray pixels) (two tailed t test, p<0.01), numbers in the top right corner indicate the coverage ratio of activated regions in V4. Red star: estimated foveal location. B. Color maps and corresponding activated regions for different spatial frequencies, acquired from the same case as in panel A. Gratings above the maps indicate the subtraction pair for the maps. Left panel: differential maps in response to R/G gratings (corresponding to the black patches) versus W/B gratings (corresponding to the white patches). Right panel: activated color preference regions for the stimuli, only including color pixels showing significantly stronger responses to R/G gratings (two tailed t test, p<0.01). C. Selective activation centers of the activated orientation domains (blue dots) and color domains (orange dots) under different SF conditions. The dots are the geometric centroid of all corresponding activated regions in V4 (two tailed t test, p<0.01). The values indicate the SF used. D, E. Combined results generated by superimposing pixels in A and B, respectively. Scale bar, 1 mm.

**Figure 5.**
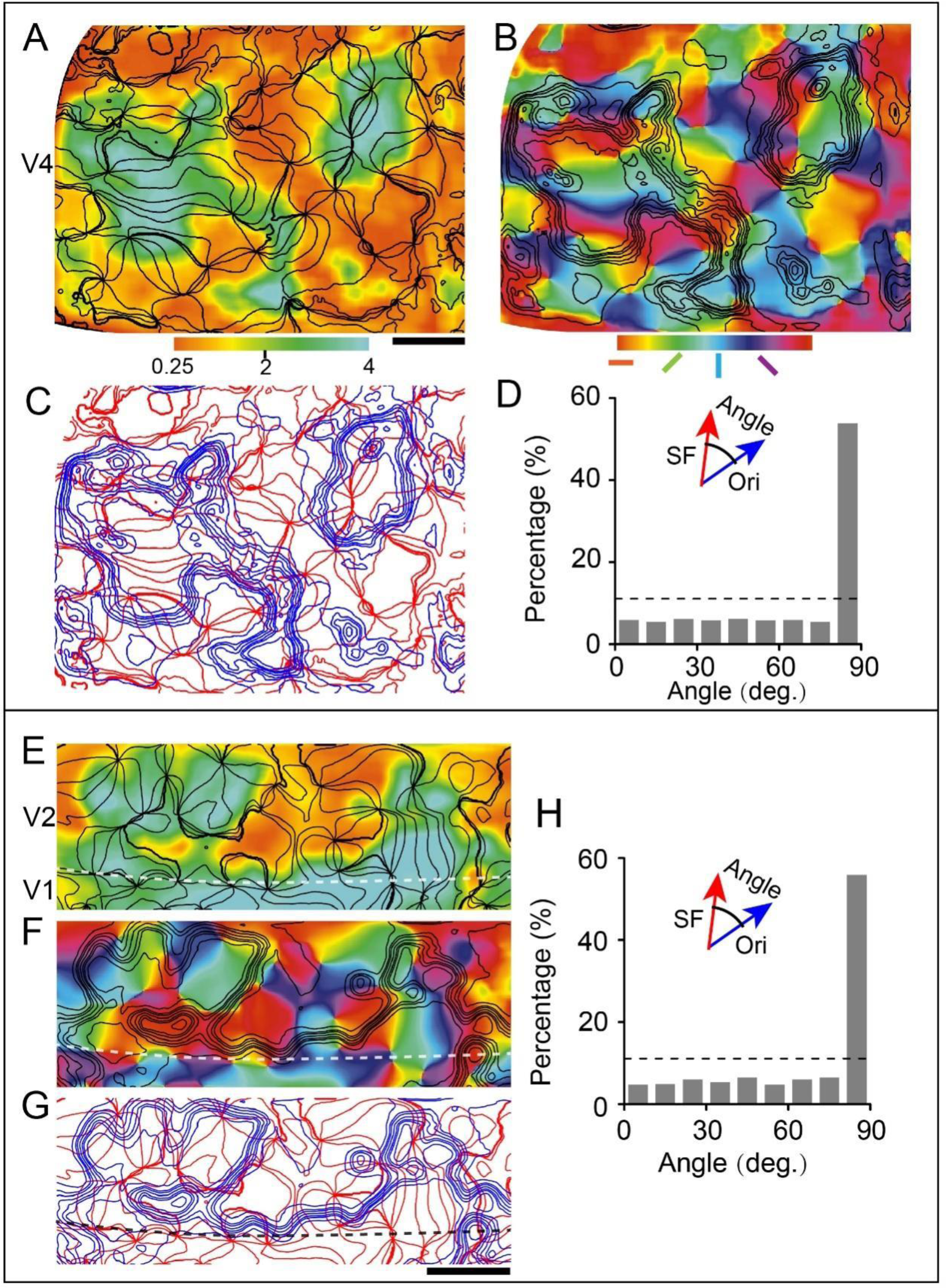
Relationship between V4/V2 spatial frequency and orientation maps. Same case as in Figure 3B, the region outlined by black dashed lines in the lateral part were chosen for this analysis (A-D, V4 region on the lateral; E-H, V2 region). A, E. Spatial frequency preference maps with the smoothed iso-orientation contours overlaid in black. B, F. Orientation angle preference maps with the smoothed iso-spatial frequency contours overlaid in black. C, G. Combined orientation (red) and spatial frequency (blue) contours. D, H. Histograms showing the distribution of the intersection angle between orientation (Ori) and spatial frequency (SF) map gradients.

### SF bias in V4 color domains

Since the first report of V4 SF preference domains (Lu et al., 2018), the relationship between V4 SF domains and V4 color domains remain unclear. It should be noted that SF preference *domains* are distinct from SF preference *maps*, as the domains are determined by statistical analysis and effectively distinguish high SF from low SF. Here, by comparing cortical responses recorded using high SF stimuli at two orientations: 45°, 135°, and low SF stimuli (at the same two orientations), we generated differential SF maps (see Figure 6A). A distinct segregation of light and dark regions is visible in this functional map. The black patches are the regions that prefer high SF to low SF, while the white patches have the opposite preference.

**Figure 6.**
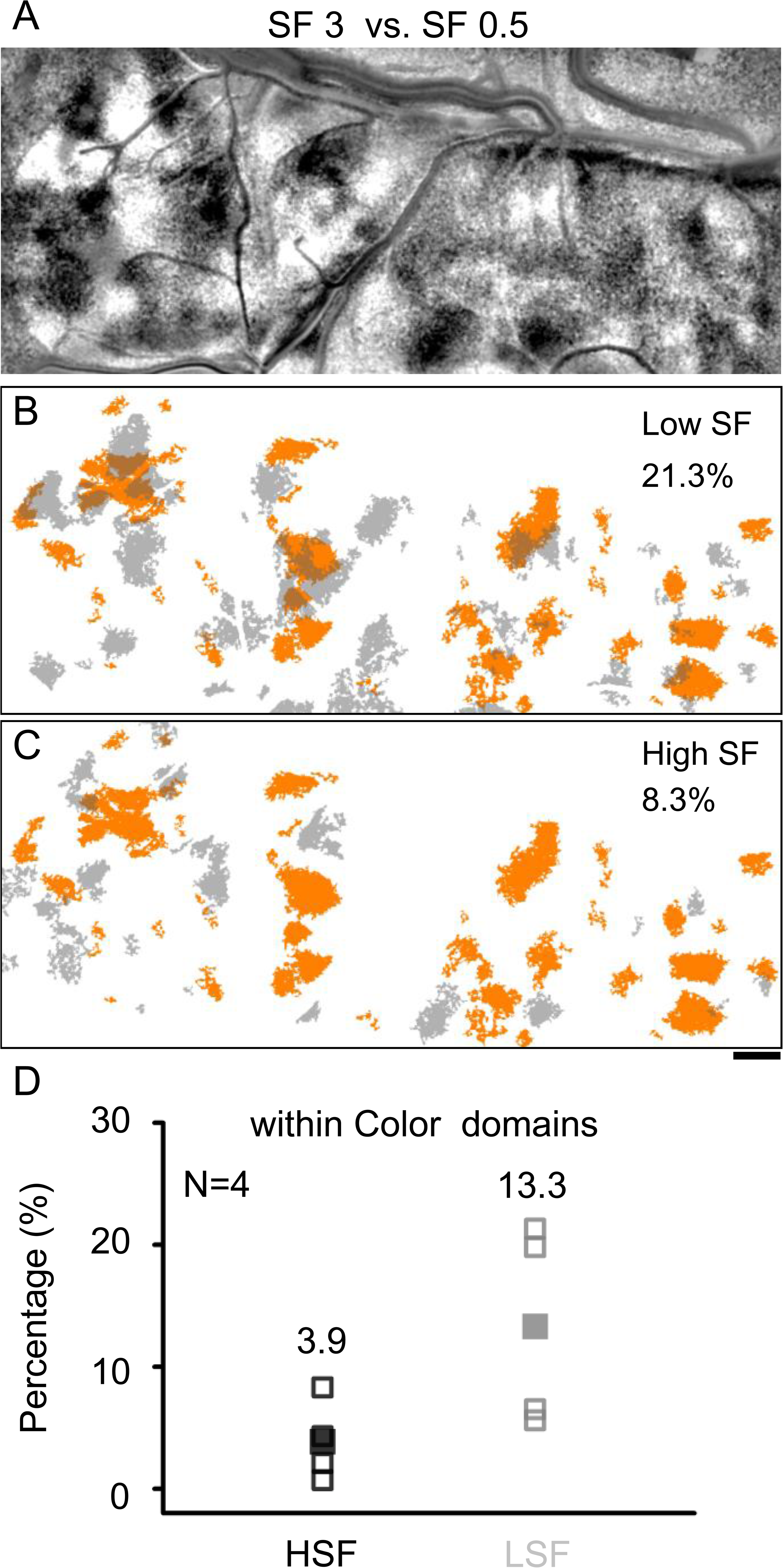
**Relationship between V4 spatial frequency and color selective domains.** A. Same case as in Figure 4A. Differential SF map in V4 is produced by subtracting the average image of two oriented grating stimuli at a low SF (0.5 cycles/deg.) from the corresponding average image at a high SF (3 cycles/deg.). The dark patches correspond to regions that prefer higher SF, while the white patches prefer lower SF. B, C. Overlay of color domains (orange) and SF domains (gray). B: low spatial frequency domains, C: high spatial frequency domains. Scale bar, 1 mm. D. The percentage of HSF/LSF selectivity regions within color domains were calculated. Unfilled squares represent the results from each case (four cases). Filled squares are the averaged outcomes from the four cases. The mean value is shown on top of the corresponding group. For other three cases, see Figure 6-figure supplement 1.

We compared the spatial relationship of color domains with SF domains (Figure 6B and C). Figure 6B shows the low SF domains color coded in gray relative to the color domains (orange patches). Similarly, high SF domains are color coded in gray relative to the same domains in Figure 6C. In this case, low SF preference domains appear to co-localize with V4 color domains to a greater extent than high SF preference domains (21.3% vs. 8.3%).

More examples are shown in Figure 6-figure supplement 1 (three additional cases from three different hemispheres). In these four cases, we found only a small portion of V4 areas discriminate high SF from low SF (two tailed t test, p<0.01; coverage ratio of high SF preference domains in V4: 3.9%±1.5%, mean±SD.; coverage ratio of low SF preference domains in V4: 6.0%±3.7%, mean±SD.). Although the difference is not statistically significant (four cases, Wilcoxon test, p>0.05), we find in all cases there is a tendency that compared to high SF preference domains, more low SF preference domains overlap with color domains (the percentage of high SF preference regions in color domains: 3.9%±3.3%; the percentage of low SF preference regions in color domains: 13.3%±8.4%, Figure 6D).

### Periodic change of SF preference in V2

Previous research in V2 has shown that relying on spatial frequency to distinguish different types of stripes (thin, pale, thick) is difficult (Levitt et al., 1994; Lu et al., 2018; Tootell and Hamilton, 1989). However, one study has reported a stripe specificity in spatial frequency selectivity (Lu and Roe, 2007). We hypothesize that this controversy is due to the inability for cytochrome oxidase staining to reliably identify stripe type, something which functional imaging can securely address. We first checked whether there were measurable SF preference differences in V2. As indicated in the map, the SF preference changed periodically (see V2 in Figure 3B, region between lunate sulcus and V1/V2 border indicated by red arrow). To compare with previous studies, we adopted the same method as Lu (Lu et al., 2018) to obtain a differential SF map (Figure 7B). The low SF preference regions obtained using these two methods (white patches, in the differential SF map; red/orange regions, in the colored SF preference map) are well correlated with color preference domains (yellow/red dashed circles in Figure 7A and B) In addition, we acquired differential SF maps by subtractions between different SF pairs (see Figure 7-figure supplement 1). For subtraction between high SF (>1 cycles/deg.) and low SF (0.5 cycles/deg.) conditions, high SF preference domains can be detected in the foveal region (red star, see black patches indicated by red arrows in Figure 7-figure supplement 1A and B). However, for a subtraction between medium SF (1 cycles/deg.) and low SF (0.25 cycles/deg.) conditions, high SF preference domains can only be detected in the parafoveal region (see black patches indicated by blue arrows in Figure 7-figure supplement 1C). These results indicate that in V2 parafoveal regions, high vs low SF preference remains stable across different SF pairs. and from parafoveal to foveal region, the preferred SF tends to gradually increase.

**Figure 7.**
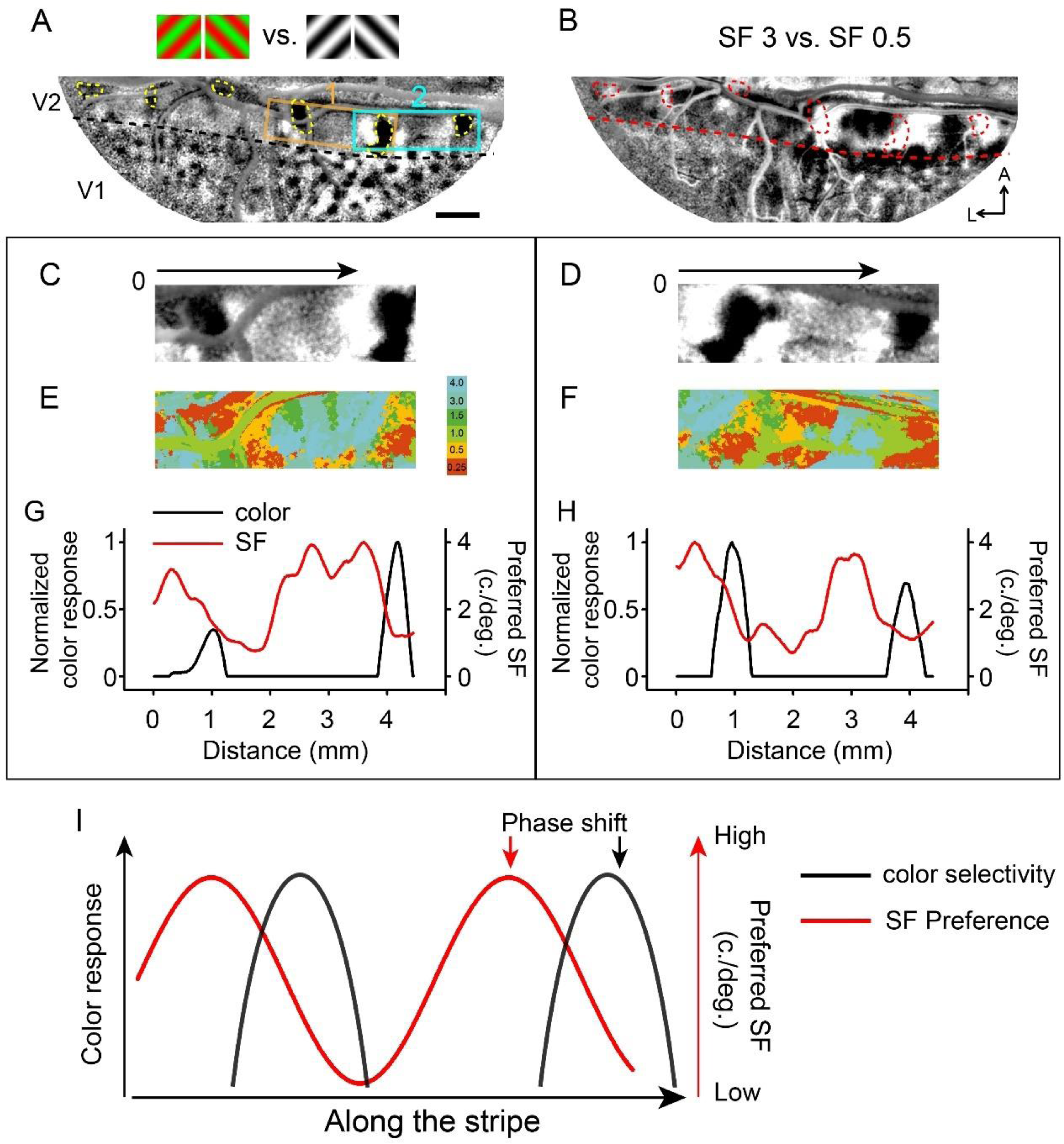
**Stripe-like distribution of SF preference in V2.** Evidence of stripe-like distribution of SF preferences in V2. Data from the same case as shown in Figure 3, Case 1. A. Color map. Regions 1 (orange rectangle) and 2 (cyan rectangle) were selected for further analysis in C-H. In V2, the yellow dashed circles highlight the color domains. The border between V1 and V2 is indicated by a black dashed line. Scale bar, 1 mm. B. Differential SF maps produced by subtracting the average image of two oriented grating stimuli at a low SF (0.5 cycles/deg.) from the corresponding average image at a high SF (3 cycles/deg.). Red dashed circles: color domains same with in (A). C, D. Enlarged color maps from regions 1 and 2. E, F. Enlarged SF maps from regions 1 and 2. G, H. Changes of color selective response (black lines) and SF preference (red lines) along the path parallel to V1/V2 border in V2. I. Similar to color selective responses in V2, SF preference changes periodically, but with a different phase.

To provide an additional illustration of how SF preference varies along V2, we selected V2 regions that exhibited two separated color domains (Figure 7A, Regions 1 and 2) and compared the changes of SF preference against color selectivity along the V2 long axis (parallel to the black arrows in Figure 7C and D). Both SF preference (red lines) and color selectivity (black lines) vary periodically (Figure 7C-H). Color selectivity was found in or near low SF preference regions (see also Figure 7B, red dashed circles and white patches in V2); however, not all low SF preference regions exhibited strong color selectivity. Thus, there are SF preference differences in V2 that vary according to a unique periodicity related to stripe periodicity (see the difference in phase between SF and color selective responses in Figure 7I).

## Discussion

Using optical imaging of intrinsic signals in a large area of visual cortex, we were able to simultaneously characterize the functional architecture underlying color, orientation, and spatial frequency domains in areas V2 and V4. We found (1) *Topography*: With respect to topography, the SF population response of V2 and V4 orientation and color domains shifts systematically with topographic location. To be specific, the geometric centroids of the selective response shift towards the lateral part of the cortex (foveolar cortex) as stimulus SF increases. This finding quantifies, at a population level, the gradation of SF preference across the visual field. Interestingly, distinct from V1, which is selectively responsive only to high SF, the foveal region of V4 is responsive to a broad range of SF from high to low, suggesting greater SF integration at higher cortical levels, even in foveal regions. (2) *Orthogonal primary axes*: Within V4 and V2, similar to what was previously shown in V1, the gradients of orientation and SF maps are predominantly orthogonal to one another. However, within color domains, which overlaps only with low SF preference domains, there is no clear gradient of SF representation. (3) *Periodicity*: We find a periodicity of spatial frequency preference fluctuation, most clearly seen in V2 but also present in V4. Specifically, this is the first illustration of a SF periodicity in relation to the V2 stripe cycle.

### General organization rules of cortical space

As mentioned in many previous studies, orthogonal crossings between different cortical maps facilitate the maximum combination of response properties in a local area with columnar organization. This kind of spatial relationship has now been reported in different animals (cat, monkey) and between different functional maps including: orientation vs. ocular dominance, orientation vs. SF in cat area 17 (Hübener et al., 1997); orientation vs. ocular dominance, orientation vs. SF in monkey V1 (Bartfeld and Grinvald, 1992; Nauhaus et al., 2012; Obermayer and Blasdel, 1993); hue vs. lightness in macaque V1 (Li et al., 2022); orientation vs. disparity in macaque V2 (Chen et al 2008; Ts’o et al., 2009); orientation vs. SF in macaque V2 and V4 (this study). These examples suggest that, to effectively use cortical space, this orthogonality is established by pairs of key visual attributes.

That is, for object structure, orientation and spatial frequency are two key parameters; for color, key parameters are hue and luminance (Li et al., 2022). These combinations ensure a complete representation of basic shape and surface information at each cortical level. Thus, cortical mosaics contain distinct regions of orthogonal feature parameters, as observed in the color and orientation stripes of V2 and the color and orientation bands of V4.

### Population selective responses across cortical space

Cortical space in its natural state includes the coding strategy and distinguishes one visual area from another visual area (V1 vs. V4) or one species from another species (monkey vs. cat). In this study, we looked at the cortical space from two perspectives: population selective response and configurational changes. Although several studies have shown how responses of functional domains changes as a result of different SFs (Issa et al., 2000; Lu and Roe, 2007; Lu et al., 2018), these studies have primarily focused on response amplitude changes. Instead of measuring the amplitudes, we characterized configurational changes in cortical space, which reflects a different measurement of cortical activity. Consistent with previous findings (Desimone and Schein, 1987; Lu et al., 2018) our results indicate that the majority of V4 regions favor low SF (Figure 3). For both V4 orientation and color selective response, there is a tendency for increasing SF to cause a shift from parafovea to fovea (Figures 4A, B). These findings support previous research that found an inverse relationship between SF preference and retinal eccentricity (Desimone and Schein, 1987; Lu et al., 2018). However, it should be noted that in V4 lateral region, even at low SF robust orientation and color selective responses are detected. The difference between V1 and V4 at fovea also indicates that V4 is better organized than V1 for processing complex images (natural scenes) with multiple SF components.

### Periodic SF preference change in V2

We explored whether SF is spatially organized similarly to other attributes in V2. Comparable to a previous study (Lu and Roe, 2007), we detected a stripe-like SF preference distribution in V2 (Figure 7). Rather than comparing this distribution to CO staining patterns, we analyzed the spatial relationship of these SF preference regions with color domains. We found that similar to color selective response, SF preference changes periodically along V2. However, we find that this repetition cycle is slightly phase shifted relative to the color cycle (Figure 7), potentially due to activation in the color-associated pale stripes (Felleman et al., 1997). This phase shift could underlie the controversy underlying CO stripe type (thin, thick, pale) and SF preference.

Why does SF preference in V2 change in this way? As suggested by modelling V2 retinotopic maps of tree shrew (Sedigh-Sarvestani et al., 2021), in elongated cortical areas such as V2 there tend to be periodic changes of response features across the cortical surface. One putative functional implication of this periodic distribution is to aid in the integration of various information (wiring minimization) across different domains via horizontal connections in V2 (Baldwin et al., 2012; Lund et al., 1993) which requires further studies.

### Thoughts about the hypercolumn

Based on the above findings, we suggest a nested hierarchy of organizations (see Figure 1). At the scale of visual field representation, there is a broad and downward shifting range of SF’s from center to periphery. Within this global SF map, lies hypercolumns of repeated orientation and color representation, each of which contains two orthogonally arranged primary parameters. In the ‘orientation’ regions, SF is systematically and orthogonally mapped in relation to the orientation map (this study); in the ‘color’ regions, hue and lightness are orthogonally mapped (Li et al., 2022). Analogous to how the primary parameter spaces mapped in each cortical area change from one area to another (e.g. V1: ocular dominance, orientation, color; V2: higher order orientation, hue, disparity; V4: curvature, hue & lightness; face areas: face maps, Kanwisher et al., 1997, Chang and Tsao, 2017; object areas: object maps, Bao et al., 2020), we hypothesize that subregions of a cortical hypercolumn also organize for different parameters. Thus, while SF is an important primary axis in orientation regions, in color regions which by nature are associated with low SF’s, the rationale for a systematic SF map is weakened. In fact, one could view the color regions as evolutionary ‘add-ons’ which became tacked on to the low end of the SF continuum. We suggest the architecture of SF representation, which describes distinct SF representations within the orientation and color regions, further extends and supports the view of a continued parallel streaming of feature-specific pathways.

## Materials and Methods

Data was acquired from 5 hemispheres of 3 adult macaque monkeys (one male and two female, Macaca mulatta). All procedures were performed in accordance with the National Institutes of Health Guidelines and were approved by the Zhejiang University Institutional Animal Care and Use Committee (Approval No. ZJU20200022 and ZJU20200023).

### Animal preparation

Chronic optical windows were implanted in contact with cortex above areas V1, V2, and V4, containing lunate sulcus and superior temporal sulcus as described previously (Hu et al., 2020, also see Figure 2B). The eccentricity of the visual field corresponding to exposed V4 was 0-5° and for V1/V2 was 0-2°. Following the craniotomy surgery, optical images were acquired in order to generate basic functional maps. Monkeys were artificially ventilated and anesthetized with propofol (induction 5-10 mg/kg, maintenance 5-10 mg/kg/hr., i.v.) and isoflurane (0.2-1.5%). Anesthetic depth was assessed continuously by monitoring heart rate, end-tidal CO2, blood oximetry, and EEG. Rectal temperature was maintained at 38 °C. Animals were paralyzed (vecuronium bromide, induction 0.25 mg/kg, maintenance 0.05-0.1 mg/kg/hr., i.v.) and respirated. Pupils were dilated (atropine sulfate 1%) and eyes fitted with contact lenses of appropriate curvature to focus on a stimulus screen 57 cm from the eyes.

### Visual stimuli for optical imaging

Visual stimuli were created using ViSaGe (Cambridge Research Systems Ltd.) and displayed on a calibrated 27-inch monitor (Philips 272G5D) operating at 60 Hz refresh rate. The luminance for white stimuli was 206.52 cd/m^2^ and black was 0.50 cd/m^2^. Full-Screen visual grating stimuli were used to locate different functional domains. To acquire color maps (Figures 4B, 7A and S2), red/green isoluminance and black-white sine-wave drifting grating stimuli, as shown in Figure 2A, were presented at two different orientations (e.g. 45° and 135°) with various spatial frequencies. To acquire orientation maps (differential orientation maps: Figures 4A and S1; orientation preference angle maps: Figures 5B, 5F, S4F and S4J) and spatial frequency maps (differential SF maps: Figures 6A, 7B; SF preference maps: Figures 3B, 3E, 5A, 5E, S4E and S4I), gratings with different orientations (0°, 45°, 90°, 135°) and different SFs (0.25, 0.5, 1, 1.5, 3, 4 cycles/deg.) were presented.

### Optical Imaging

The brain was imaged through a glass window mounted in contact with cortex. Images of cortical reflectance changes (intrinsic hemodynamic signals) corresponding to local cortical activity were acquired (Imager 3001, Optical Imaging Inc., German town, NY) with 630 nm illumination. Image size was 1080 ×1308 pixels representing 14.4×17.4 or 8.7×10.5 mm field of view. Visual stimuli were presented in a random order. Each stimulus was presented for 3.5-4.5 seconds. Frames were acquired at 4 Hz for 4-5 seconds synchronized to respiration. Visual stimuli were presented 0.5 seconds after beginning image acquisition. The imaging data were stored in a block fashion. Each block contained the imaging data recorded from the stimulus conditions (presented one time). Each stimulus was presented at least 25 times.

### Data analysis

#### Generation of functional maps

With the following formula, 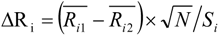, we assessed the response differences between two comparison groups. 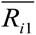 and 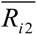 are the mean dR/R’ values (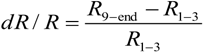, R9-end is the averaged response from frames 9 to the last frame, R1-3 is the averaged response from frames 1 to 3) in the two compared conditions for pixel i, N is the number of trials, and Si is the standard deviation of (*R_i_*_1_ − *R_i_* _2_). Single condition maps were obtained by comparing the images acquired during stimulus and during a blank.

Color maps were obtained by comparing red/green and white/black grating images, differential spatial frequency maps were obtained by comparing high (2-6 cycles/deg.) and low (0.25-0.5 cycles/deg.) spatial frequency images, and differential orientation maps were obtained by comparing two orthogonal orientation images (45° vs. 135°). Maps were low-pass filtered (Gaussian filter, ∼30-80 μm diameter) and low-frequency noise was reduced by convolving a given map with a ∼1-2 mm diameter Gaussian filter and subtracting from the original map. Within a single experimental session, the same filtering parameters were always used to ensure that this filtering procedure did not influence the observed differences. The border between V1 and V2 was determined based on color map: in V1 color response has a blob-like distribution whereas in V2 color response has a stripe-like distribution.

To generate spatial frequency preference (SFP) maps (e.g. in Figure 3B, 3E), for each pixel we compared its activation under different single SF conditions. The preferred SF of each pixel was defined as the SF at which the strongest activation signal for that pixel was observed. The comparison includes 2 orientations, (45° and 135°); for each orientation, six different SFs, 0.25, 0.5, 1, 1.5, 3, 4 cycles/deg. were presented.). Each pixel in a given SFP map was assigned a unique color to represent the preferred SF. Orientation angle preference maps (Figures 5B and 5F) were calculated based on single orientation condition maps (four orientations, 0°, 45°, 90°, 135°), and each pixel was assigned a unique color to represent the preferred orientation (Bosking et al., 1997).

#### Locating the positions of selective activation and determining the activation center

Functional domains were identified by selecting the pixels with a significant difference in dR/R (two-tailed t test, p<0.01) under two comparison conditions (see Table 1).

**Table 1.**
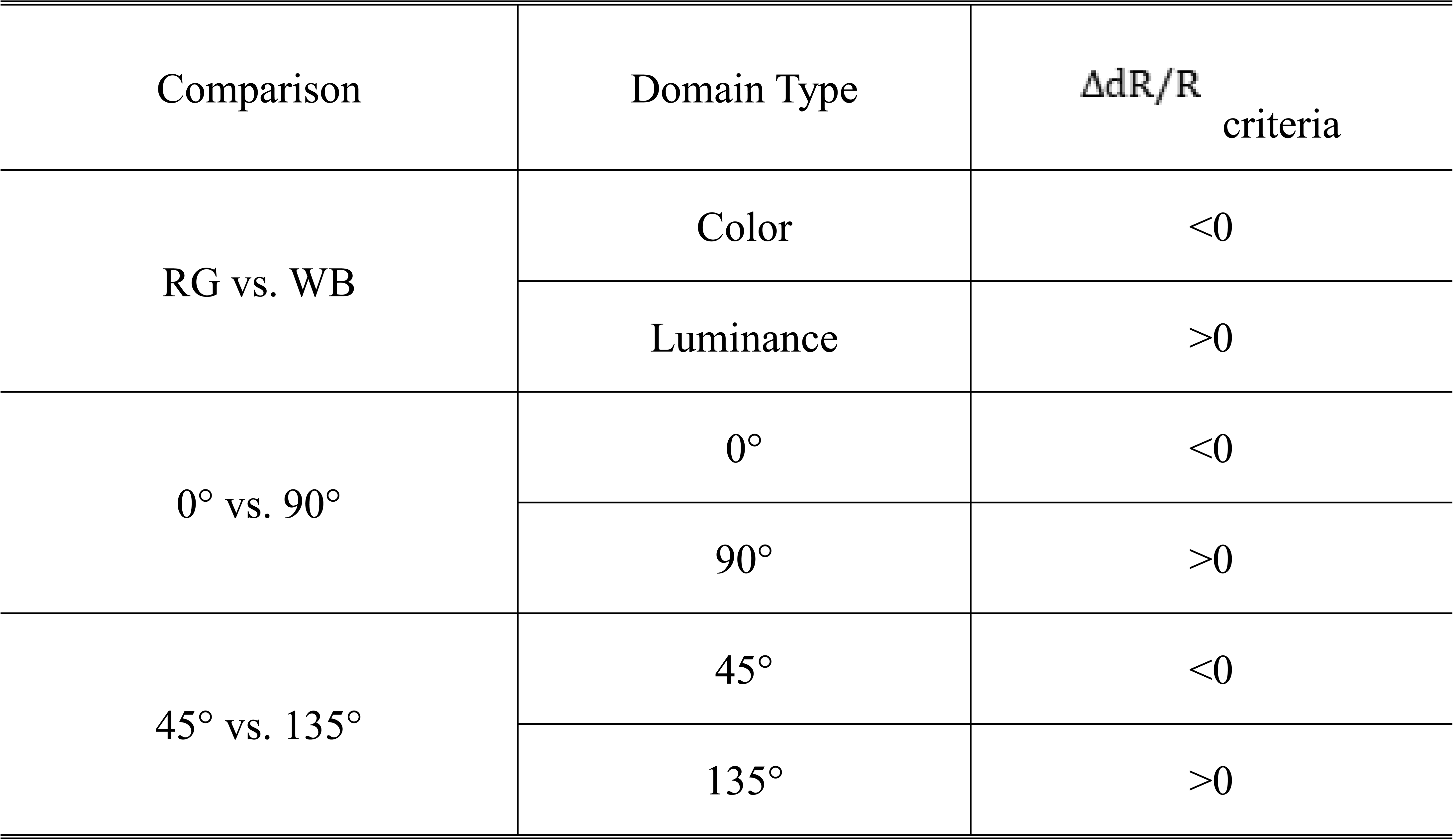
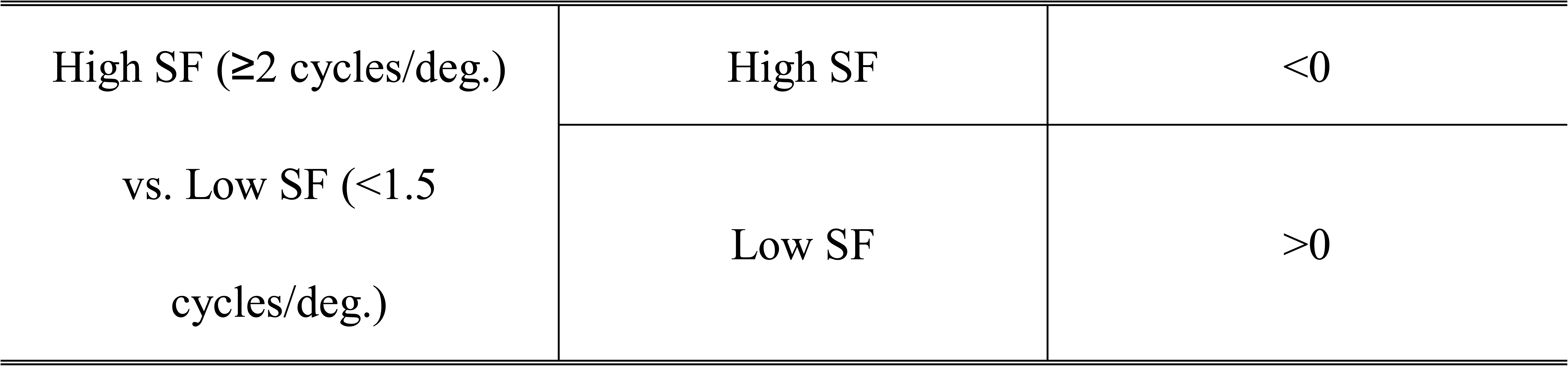
Comparisons used to generate different functional domains.

For a given activated region, the activation center was defined as the geometric centroid of all significantly activated pixels within the region; e.g., the orientation selective activation center in Figure 4A is the centroid of all the pixels of 45° and 135° orientation domains under one SF condition and the color activation center in Figure 4B is the centroid of all the pixels of color domains under one SF condition. The overlap between different functional domains was also calculated based on these thus-defined functional domains (see Figure 6).

#### Calculating the correlation of pairs of maps

To quantify the correlation between two functional maps, we isolated the significant responses in the imaged area of V4 (regions that were significantly activated by the visual stimuli, two tailed t test, p<0.01) and calculated the correlation coefficient values between the maps (Figure 4A, B, right panels) acquired under different SF conditions. To compare the difference between foveal and parafoveal regions, we divided the imaged V4 regions into two halves: the left half of the region was designated as foveal, the right half was designated as parafoveal, and the correlation value for each half was calculated separately.

#### Comparing the spatial relationship between SF and orientation maps

The spatial frequency preference maps and orientation angle preference maps were smoothed (high-pass filtered with a Gaussian filter, ∼200 μm diameter) before obtaining contours and gradients. Iso-orientation contours (8 contours, 0°, 22.5°, 45°, 67.5°, 90°, 112.5°, 135°, 157.5°) and iso-SF contours (5 contours, 0.25, 0.5, 1, 2, 4 cycles/deg.) were drawn based on these smoothed maps using the MATLAB ‘contour’ function. The gradient directions of the smoothed SF and orientation maps were calculated at each pixel using the MATLAB ‘gradient’ function; the spacing used for gradient calculations in each direction was ∼50 μm. The difference between the two gradients was calculated to determine the spatial relationship between SF and orientation maps (e.g. Figure 5D, H).

#### Characterizing the periodicity of SF preference in V2

As reported in previous studies, V2 color selective response changes periodically along the long axis of V2 (Levitt et al., 1994; Roe and Ts’o, 1995). To characterize the periodic change of SF preference in V2, we chose a region of V2 with clearly identifiable periodic changes in color response (at least two well-separated color domains) for further analysis. We slightly rotated the selected V2 region in order to align the V1/V2 border horizontally in the cropped small map (see Figure 7C and D). For each of these small maps, we quantified and normalized color selectivity and SF preference for all pixels in the map. The average value for the pixels along each vertical line at different distances from left (distance = 0 mm) to right (distance = 4.5 mm) were then computed and plotted.

## Acknowledgements

We thank Yin Liu, Meizhen Qian for help with the animal experiments. This research was conducted at Zhejiang University and was supported by the National Natural Science Foundation of China (grant nos. U1909205, U20A20221, and 81961128029 to A.W.R.), the National Key R&D Program of China (grant no. 2018YFA0701400 to A.W.R.), Chinese NSF Instrumentation (grant no. 31627802 to A.W.R.), Key Research and Development Program of Zhejiang Province (grant no. 2020C03004 to A.W.R.), the Fundamental Research Funds for the Central Universities (grant no. 2019XZZX003-20 to A.W.R.), China Postdoctoral Science Foundation (grant no. 2020M681829 to J.M.H.), National Natural Science Foundation of China (grant no. 32100802 to J.M.H.), and the MOE Frontier Science Center for Brain Science & Brain-Machine Integration, Zhejiang University.

---

**Figure 4—figure supplement 1.**
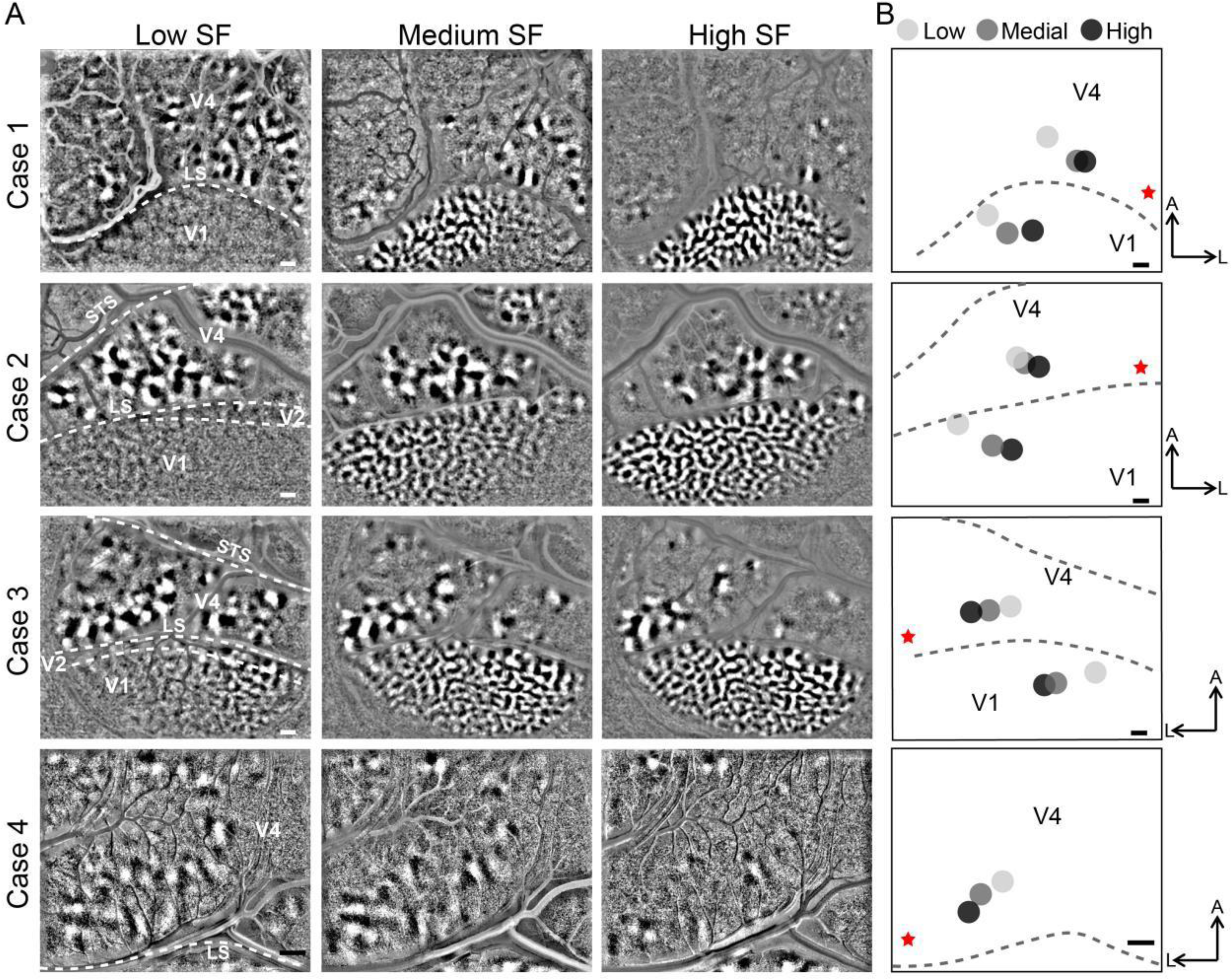
Orientation maps obtained by using drifting gratings of different SFs. A. Imaging results of cortical responses to gratings with different spatial frequencies (indicated on top in each column). Maps generated from response difference of 45° gratings minus 135° gratings (each row represents results from different cases). LS, lunate sulcus; STS, superior temporal sulcus. B. Spatial distribution of the geometric center of activated orientation domains at different SFs. Under different SF conditions, the geometric centers of the activated orientation domains were calculated (two tailed t test, p<0.01). Light gray: low SF, 0.25-0.5 cycles/deg; gray: medium SF, 1-2 cycles/deg; dark gray: high SF, 3-4 cycles/deg. V1 and V4 activation centers were labeled separately. L, lateral; A, anterior; LS, lunate sulcus; Red star, estimated foveal location. Scale bar, 1mm.

**Figure 4—figure supplement 2.**
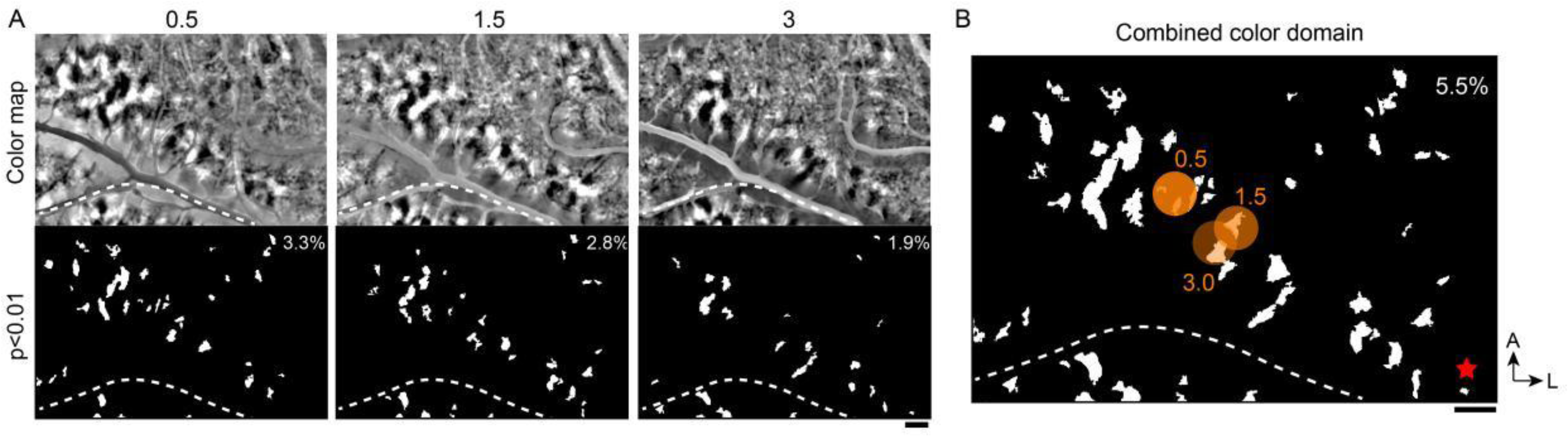
Another example of V4 color map acquired with different SFs. A. Color maps and corresponding activated regions in different spatial frequencies. Top panel, differential maps in response to R/G gratings (corresponding to the black patches) versus W/B gratings (corresponding to the white patches); bottom panel, activated regions by the stimuli, only including color pixels (show significantly stronger responses to R/G gratings, two tailed t test, p<0.01). B. Combined results from A. Pixels in A are superimposed. Numbers in the upper right corner represent the coverage ratio of activated color domains in V4. Orange dots: centers of activated color domains corresponding to different SFs. Red star: estimated foveal location. Scale bar, 1mm.

**Figure 4—figure supplement 3.**
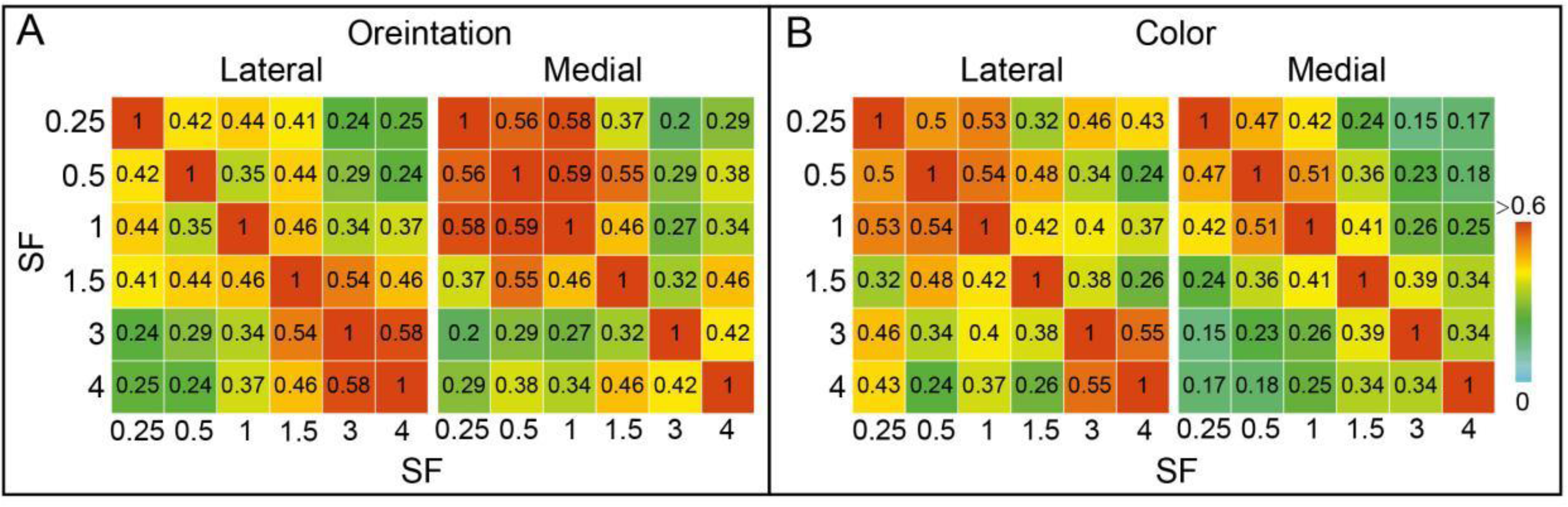
The matrices show the correlation values for pairs of maps acquired at different SFs. Same case as in Figure 4. A. Matrices of correlation values of orientation maps. B. Matrices of correlation values of color maps. The imaged V4 regions were divided into two halves: the left half of the region was designated as Lateral, the right half was designated as Medial, and the correlation value for each half was calculated separately.

**Figure 5—figure supplement 1.**
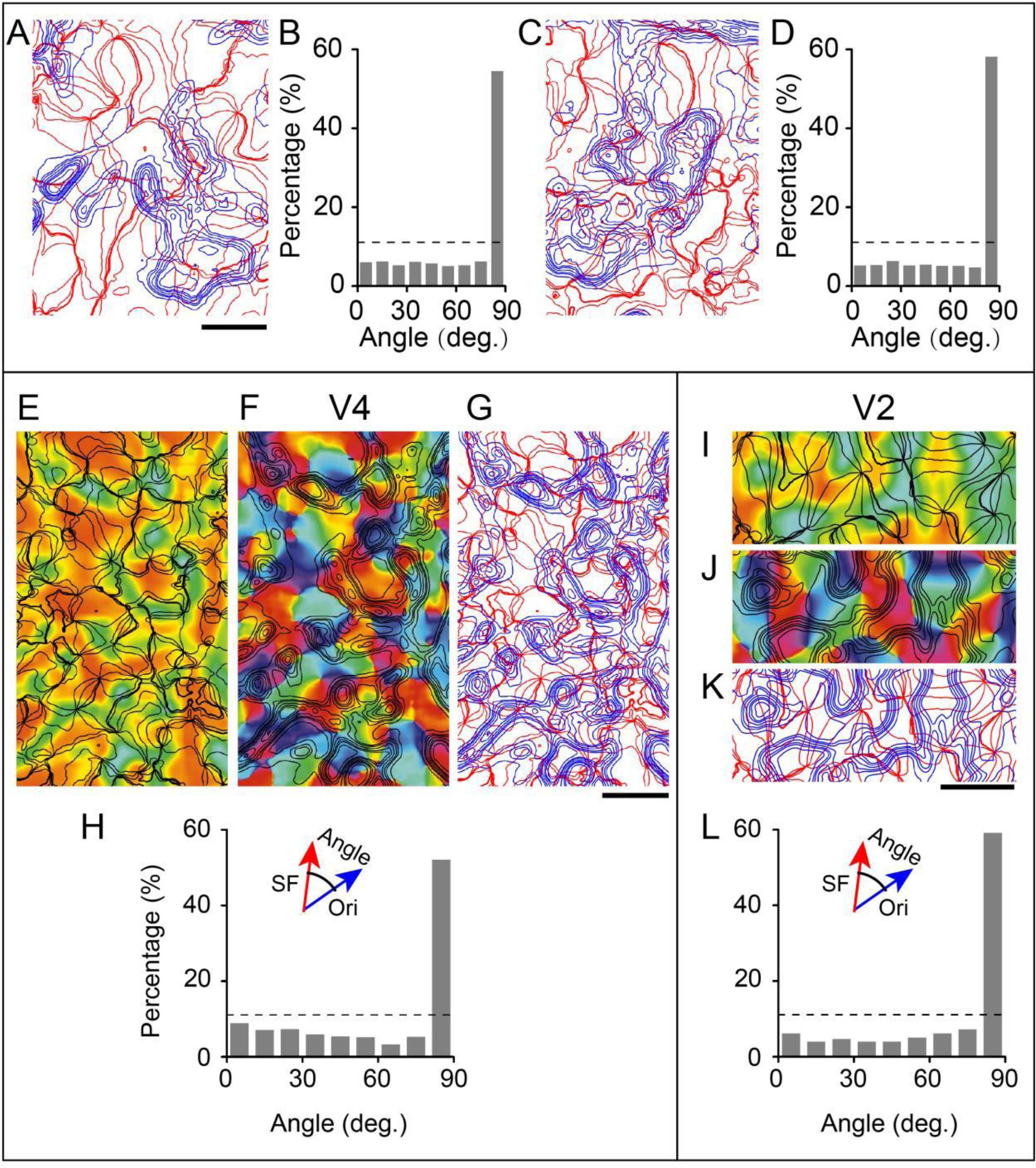
Additional examples of relationship between V4 (A-G) and V2 (I-L) spatial frequency and orientation maps. A-D. same case as in Figure 3B; E-L. same case as in Figure 3E. A, C, G, K. Combined iso-orientation (red) and iso-spatial frequency (blue) contours. E, I. Spatial frequency preference maps with the smoothed iso-orientation contours overlaid in black. F, J. Orientation angle preference maps with the smoothed iso-spatial frequency contours overlaid in black. B, D, H, L. Histograms showing the distribution of the intersection angle between orientation (Ori) and spatial frequency (SF) map gradients.

**Figure 6—figure supplement 1.**
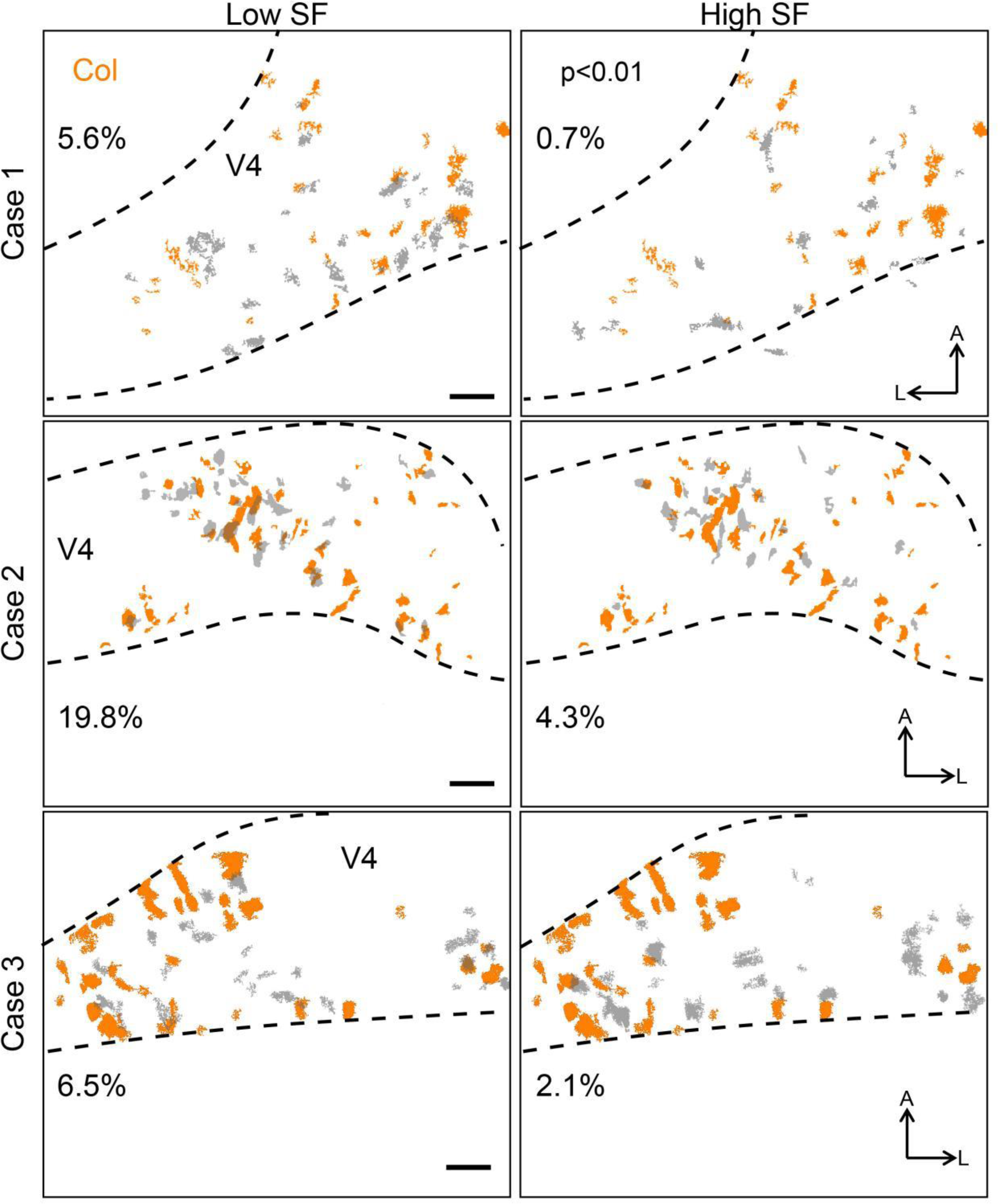
Relationships between V4 SF and color selective domains in three cases. SF domains (marked by gray) were the areas with a significant response difference (two tailed t test, p<0.01) between low SF conditions (0.25-0.5 cycles/deg, 45° and 135°, left panel) and high SF conditions (3-6 cycles/deg, 45° and 135°, right panel). Color domains (marked by orange) were superimposed over results based on activated regions (in gray) under different spatial frequency conditions. Each row of subplots represents results from different cases. Scale bar: 1mm.

**Figure 7—figure supplement 1.**
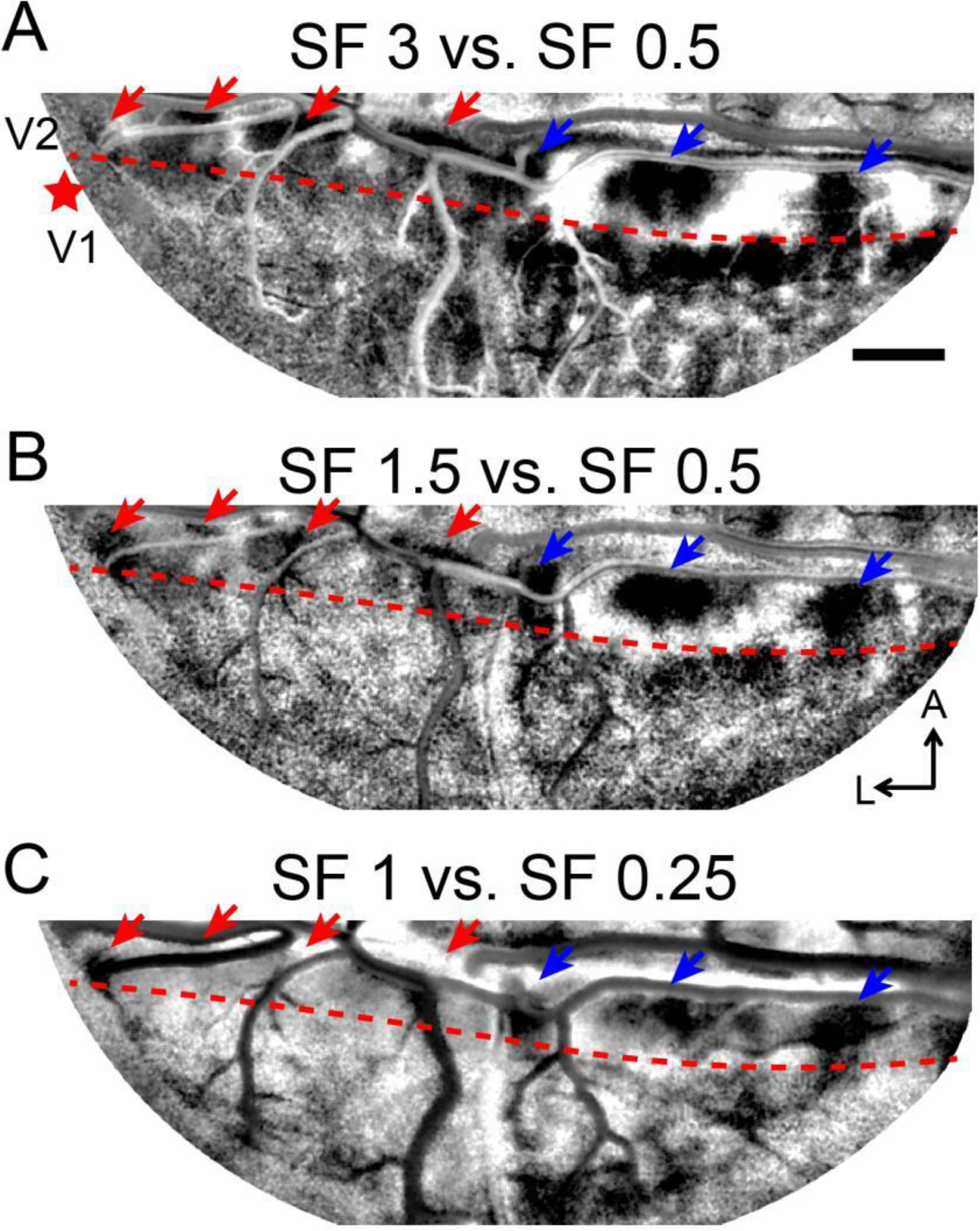
Differential SF maps acquired by subtractions between different SF pairs (same case as. Figure 7). A-C. Differential SF maps produced by subtracting the average image of two oriented grating stimuli at a low SF (A, B: 0.5 cycles/deg; C: 0.25 cycles/deg) from the corresponding average image at a higher SF (A: 3 cycles/deg; B: 1.5 cycles/deg; C: 1 cycles/deg). Red star: estimated foveal location. The arrows indicate the high SF preference regions in V2.

## References

Baldwin, M. K. L., Kaskan, P. M., Zhang, B., Chino, Y. M., & Kaas, J. H. (2012). Cortical and subcortical connections of V1 and V2 in early postnatal macaque monkeys. Journal of Comparative Neurology, 520(3), 544–569.

Bao, P., She, L., McGill, M., & Tsao, D.Y. (2020). A map of object space in primate inferotemporal cortex. Nature. 583(7814),103–108.

Bartfeld, E., & Grinvald, A. (1992). Relationships between orientation- preference pinwheels, cytochrome oxidase blobs, and ocular- dominance columns in primate striate cortex. Proc. Natl. Acad. Sci. U.S.A., 89(24), 11905–11909.

Bosking, W. H., Zhang, Y., Schofield, B., & Fitzpatrick, D. (1997). Orientation selectivity and the arrangement of horizontal connections in tree shrew striate cortex. Journal of Neuroscience, 17(6), 2112–2127.

Chang, L., & Tsao, D.Y. (2017). The Code for Facial Identity in the Primate Brain. Cell. 169(6),1013–1028.e14.

Chen, M., Li, P., Zhu, S., Han, C., Xu, H., Fang, Y., Hu, J., Roe, A. W., & Lu, H. D. (2016). An Orientation Map for Motion Boundaries in Macaque V2. Cerebral cortex, 26(1), 279–287.

Chen, G., Lu, H. D., & Roe, A. W. (2008). A map for horizontal disparity in monkey V2. Neuron, 58(3), 442–450.

Desimone, R., & Schein, S. J. (1987). Visual properties of neurons in area V4 of the macaque: Sensitivity to stimulus form. Journal of Neurophysiology, 57(3), 835–868.

Fang, Y., Chen, M., Xu, H., Li, P., Han, C., Hu, J., Zhu, S., Ma, H., & Lu, H. D. (2019). An Orientation Map for Disparity-Defined Edges in Area V4. Cerebral cortex, 29(2), 666–679.

Felleman, D. J., Xiao, Y., & McClendon, E. (1997). Modular organization of occipito-temporal pathways: cortical connections between visual area 4 and visual area 2 and posterior inferotemporal ventral area in macaque monkeys. Journal of neuroscience, 17(9), 3185–3200.

Freeman, J., Ziemba, C. M., Heeger, D. J., Simoncelli, E. P., & Movshon, J. A. (2013). A functional and perceptual signature of the second visual area in primates. Nature neuroscience, 16(7), 974–981.

Gattass, R., Gross, C. G., & Sandell, J. H. (1981). Visual topography of V2 in the macaque. Journal of Comparative Neurology, 201(4), 519–539.

Gattass, R., Sousa, A. P., & Gross, C. G. (1988). Visuotopic organization and extent of V3 and V4 of the macaque. Journal of Neuroscience, 8(6), 1831–1845.

Gegenfurtner, K. R., Kiper, D. C., & Fenstemaker, S. B. (1996). Processing of color, form, and motion in macaque area V2. Visual neuroscience, 13(1), 161–172.

Hu, J. M., Song, X. M., Wang, Q., & Roe, A. W. (2020). Curvature domains in V4 of macaque monkey. ELife, 9, e57261.

Hübener, M., Shoham, D., Grinvald, A., & Bonhoeffer, T. (1997). Spatial Relationships among Three Columnar Systems in Cat Area 17. Journal of Neuroscience, 17(23), 9270–9284.

Issa, N. P., Trepel, C., & Stryker, M. P. (2000). Spatial Frequency Maps in Cat Visual Cortex. Journal of Neuroscience, 20(22), 8504–8514.

Kanwisher, N., McDermott, J., & Chun, M. (1997) The Fusiform Face Area: A Module in Human Extrastriate Cortex Specialized for the Perception of Faces. Journal of Neuroscience. 17, 4302–4311.

Levitt, J. B., Kiper, D. C., & Movshon, J. A. (1994). Receptive fields and functional architecture of macaque V2. Journal of neurophysiology, 71(6), 2517–2542.

Li, M., Ju, N., Jiang, R., Liu, F., Jiang, H., Macknik, S., Martinez-Conde, S., & Tang, S. (2022). Perceptual hue, lightness, and chroma are represented in a multidimensional functional anatomical map in macaque V1. Progress in Neurobiology, 212, 102251.

Li, M., Liu, F., Juusola, M., & Tang, S. (2014). Perceptual color map in macaque visual area V4. Journal of Neuroscience, 34(1), 202–217.

Li, P., Zhu, S., Chen, M., Han, C., Xu, H., Hu, J., Fang, Y., & Lu, H. D. (2013). A Motion Direction Preference Map in Monkey V4. Neuron, 78(2), 376– 388.

Liu, Y., Li, M., Zhang, X., Lu, Y., Gong, H., Yin, J., Chen, Z., Qian, L., Yang, Y., Andolina, I. M., Shipp, S., Mcloughlin, N., Tang, S., & Wang, W. (2020). Hierarchical Representation for Chromatic Processing across Macaque V1, V2, and V4. Neuron, 108(3), 538–550.e5.

Lu, H. D., Chen, G., Tanigawa, H., & Roe, A. W. (2010). A motion direction map in macaque V2. Neuron, 68(5), 1002–1013.

Lu, H. D., & Roe, A. W. (2007). Optical Imaging of Contrast Response in Macaque Monkey V1 and V2. Cerebral Cortex, 17(11), 2675–2695.

Lu, Y., Yin, J., Chen, Z., Gong, H., Liu, Y., Qian, L., Li, X., Liu, R., Andolina, I. M., & Wang, W. (2018). Revealing Detail along the Visual Hierarchy: Neural Clustering Preserves Acuity from V1 to V4. Neuron, 98(2), 417–428.e3.

Lund, J. S., Yoshioka, T., & Levitt, J. B. (1993). Comparison of intrinsic connectivity in different areas of macaque monkey cerebral cortex. Cerebral Cortex, 3(2), 148–162.

Nauhaus I., Nielsen K. J., & Callaway E. M. (2016). Efficient Receptive Field Tiling in Primate V1. Neuron, 91(4), 893–904.

Nauhaus, I., Nielsen, K. J., Disney, A. A., & Callaway, E. M. (2012). Orthogonal micro-organization of orientation and spatial frequency in primate primary visual cortex. Nature Neuroscience, 15(12), 1683–1690.

Obermayer, K., & Blasdel, G. G. (1993). Geometry of orientation and ocular dominance columns in monkey striate cortex. Journal of Neuroscience, 13(10), 4114–4129.

Ponce, C. R., Hartmann, T. S., & Livingstone, M. S. (2017). End-Stopping Predicts Curvature Tuning along the Ventral Stream. Journal of neuroscience, 37(3), 648–659.

Ramsden, B. M., Hung, C. P., & Roe, A. W. (2001). Real and illusory contour processing in area V1 of the primate: a cortical balancing act. Cerebral cortex, 11(7), 648–665.

Roe, A. W., Chen, G., & Lu, H. D. (2009). Visual System: Functional Architecture of Area V2. In L. R. Squire (Ed.), Encyclopedia of Neuroscience, 331–349.

Roe, A. W., Lu, H. D., & Hung, C. P. (2005). Cortical processing of a brightness illusion. Proc. Natl. Acad. Sci. U.S.A., 102(10), 3869–3874.

Roe, A. W., & Ts’o, D. Y. (1995). Visual topography in primate V2: Multiple representation across functional stripes. Journal of Neuroscience, *15*(5 Pt 2), 3689–3715.

Sedigh-Sarvestani, M., Lee, K.-S., Jaepel, J., Satterfield, R., Shultz, N., & Fitzpatrick, D. (2021). A sinusoidal transformation of the visual field is the basis for periodic maps in area V2. Neuron, 109(24), 4068–4079.e6.

Shoham, D., Hübener, M., Schulze, S., Grinvald, A., & Bonhoeffer, T. (1997). Spatio-temporal frequency domains and their relation to cytochrome oxidase staining in cat visual cortex. Nature, 385(6616), 529–533.

Silverman, M. S., Grosof, D. H., De Valois, R. L., & Elfar, S. D. (1989). Spatial- frequency organization in primate striate cortex. Proc. Natl. Acad. Sci. U.S.A., 86(2), 711–715.

Swindale, N. V., Shoham, D., Grinvald, A., Bonhoeffer, T., & Hubener, M. (2000). Visual cortex maps are optimized for uniform coverage. Nature Neuroscience, 3(8), 822–826.

Tang, R., Song, Q., Li, Y., Zhang, R., Cai, X., & Lu, H. D. (2020). Curvature- processing domains in primate V4. ELife, 9, e57502.

Tanigawa, H., Lu, H. D., & Roe, A. W. (2010). Functional organization for color and orientation in macaque V4. Nature Neuroscience, 13(12), 1542– 1548.

Tootell, R. B., Silverman, M. S., & De Valois, R. L. (1981). Spatial frequency columns in primary visual cortex. Science, 214(4522), 813–815.

Tootell, R., & Hamilton, S. (1989). Functional anatomy of the second visual area (V2) in the macaque. Journal of Neuroscience, 9(8), 2620–2644.

Tootell, R., Silverman, M., Hamilton, S., Switkes, E., & De Valois, R. (1988). Functional anatomy of macaque striate cortex. V. Spatial frequency. Journal of Neuroscience, 8(5), 1610–1624.

Ts’o, D. Y., Zarella, M., & Burkitt, G. (2009). Whither the hypercolumn? The Journal of Physiology, 587(Pt 12), 2791–2805.

Wang, Y., Xiao, Y., & Felleman, D. J. (2007). V2 thin stripes contain spatially organized representations of achromatic luminance change. Cerebral cortex, 17(1), 116–129.

Xiao, Y., Wang, Y., & Felleman, D. J. (2003). A spatially organized representation of colour in macaque cortical area V2. Nature, 421(6922), 535–539.

Xu, X., Anderson, T. J., & Casagrande, V. A. (2007). How do functional maps in primary visual cortex vary with eccentricity? Journal of Comparative Neurology, 501(5), 741–755.

